# Development of a cross-protective common cold coronavirus vaccine

**DOI:** 10.1101/2025.05.12.653567

**Authors:** Tanushree Dangi, Shiyi Li, Pablo Penaloza-MacMaster

## Abstract

Common cold coronaviruses, such as OC43 and HKU1, generally cause mild respiratory infections in healthy people. However, they can lead to severe illness in high-risk groups, including immunocompromised individuals and older adults. Currently, there is no clinically approved vaccine to prevent infection by common cold coronaviruses. Here, we developed an mRNA vaccine expressing a stabilized spike protein derived from OC43 and tested its efficacy in different challenge models in C57BL/6 mice. This novel OC43 vaccine elicited OC43-specific immune responses, as well as cross-reactive immune response against other embecoviruses, including HKU1 and mouse hepatitis virus (MHV-A59). Interestingly, this OC43 vaccine protected mice not only against a lethal OC43 infection, but also against MHV-A59, which is only 65% matched. Vaccine cross-protection appeared to be mechanistically mediated by non-neutralizing antibodies, but not by CD8 and CD4 T cells. These findings provide insights for the development of common cold coronavirus vaccines, demonstrating the potential for a single vaccine to target different members of a coronavirus subgenus.

## INTRODUCTION

SARS-CoV-2 has resulted in over 7 million deaths globally. However, the rapid development of SARS-CoV-2 vaccines reduced the death toll of the pandemic. In addition to SARS-CoV-2, several endemic coronaviruses like OC43, 229E, HKU1, and NL63 circulate within the human population, causing frequent re-infections every year. While these endemic coronaviruses typically cause mild to moderate upper respiratory infections, they can also lead to severe conditions pneumonia and bronchitis, particularly in the elderly and immunocompromised individuals (1–6). Among these, the human common cold coronavirus, OC43, causes frequent respiratory infections worldwide, but no effective vaccine is currently available.

The widespread circulation of OC43 poses public health concerns due to its propensity for mutation and potential recombination with other coronaviruses. OC43 causes a considerable burden on healthcare systems, resulting in significant economic losses. The high incidence of OC43 reinfections further highlights the need for effective vaccines (7–10). Addressing these challenges with an effective vaccine would not only improve overall health in the population but also has the potential to alleviate the economic burden associated with recurrent coronavirus infections. This motivated us to develop an mRNA vaccine encoding a stabilized OC43 spike protein. Our results demonstrate that this vaccine is immunogenic and highly protective not only against OC43, but also a distant embecovirus.

## RESULTS

### Design of a novel mRNA-LNP vaccine encoding HCoV-OC43 spike glycoprotein (mRNA-OC43)

We developed an mRNA vaccine encoding the full-length spike glycoprotein of the OC43 coronavirus. We utilized the spike protein sequence from NCBI (AAA03055.1) and introduced two proline mutations at 1070 and 1071 amino acid positions (replaced with A and L amino acids) in the S2 domain to stabilize the spike protein in its pre-fusion state. A pcDNA3.1 (+) plasmid construct was designed by incorporating the codon-optimized spike gene sequence flanked with untranslated regions (UTRs) at 3’ and 5’ ends, and a T7 RNA polymerase site before the 3’ end of coding sequence. Expression of OC43 spike glycoprotein was confirmed by transfecting HEK293T cells with the corresponding mRNA followed by Western Blot analysis to validate protein expression (Fig. 1). mRNA molecules were then encapsulated into a lipid nanoparticle using a Nanoassemblr.

**Figure 1.**
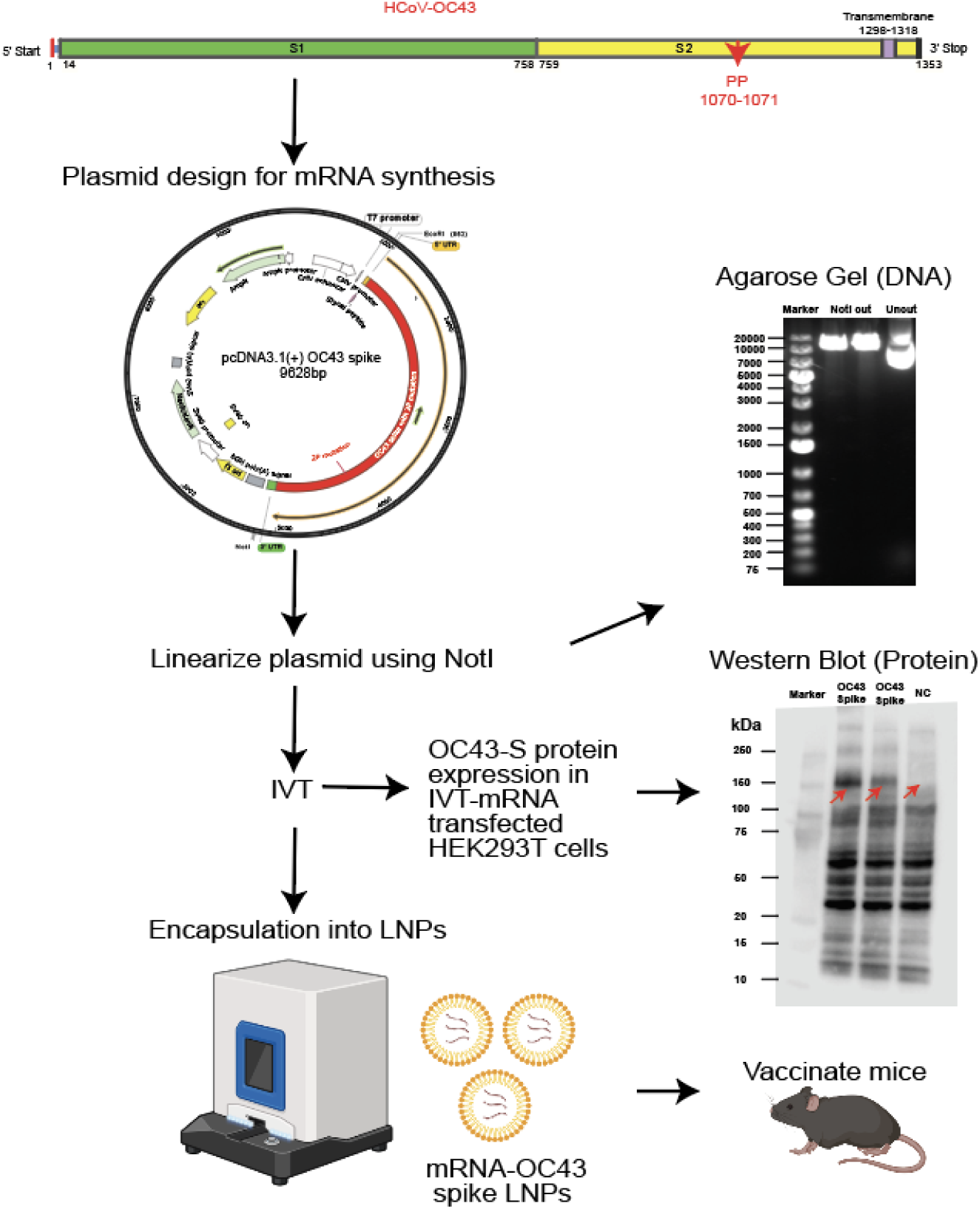
Formulation strategy of mRNA-based common cold coronavirus vaccine encoding OC43-spike protein. The spike gene sequence of the OC43 was modified by inserting two proline mutations in the S2 subdomain and a codon-optimized for mouse strain. pcDNA3.1(+) plasmid was constructed by incorporating modified OC43-spike gene. Before proceeding with the IVT reaction, the plasmid construct was linearized using NotI restriction digestion, and its size was confirmed by agarose gel electrophoresis. mRNA was transcribed in vitro (IVT-mRNA) using a plasmid construct and incorporated with 0 cap and poly-A tail at 5’ and 3’ ends of transcribed mRNA. The expression of the OC43 spike gene was tested in HEK293T cells transfected with mRNA-OC43 and by analyzing the transfected cell lysate in western blot. Vaccine was prepared by encapsulating IVT-mRNA into lipid nanoparticles utilizing the Nanoasemblr platform.

### Immunogenicity and Protective Efficacy of mRNA-OC43 following homologous OC43 challenge

To evaluate immunogenicity, we first immunized C57BL/6 mice intramuscularly with 3 μg of mRNA-OC43 (Fig. 2A). At day 15 post-vaccination, we measured antibody and T cell responses in plasma, using enzyme-linked immunosorbent assay (ELISA) with OC43 spike as coating antigen, and intracellular cytokine assay (ICS) using overlapping OC43 spike peptide pools. The mRNA-OC43 vaccine elicited potent antibody responses to the OC43 spike protein (Fig. 2B). Moreover, the vaccine elicited CD8 and CD4 T cell responses by ICS (Fig. 2C-2F).

**Figure 2.**
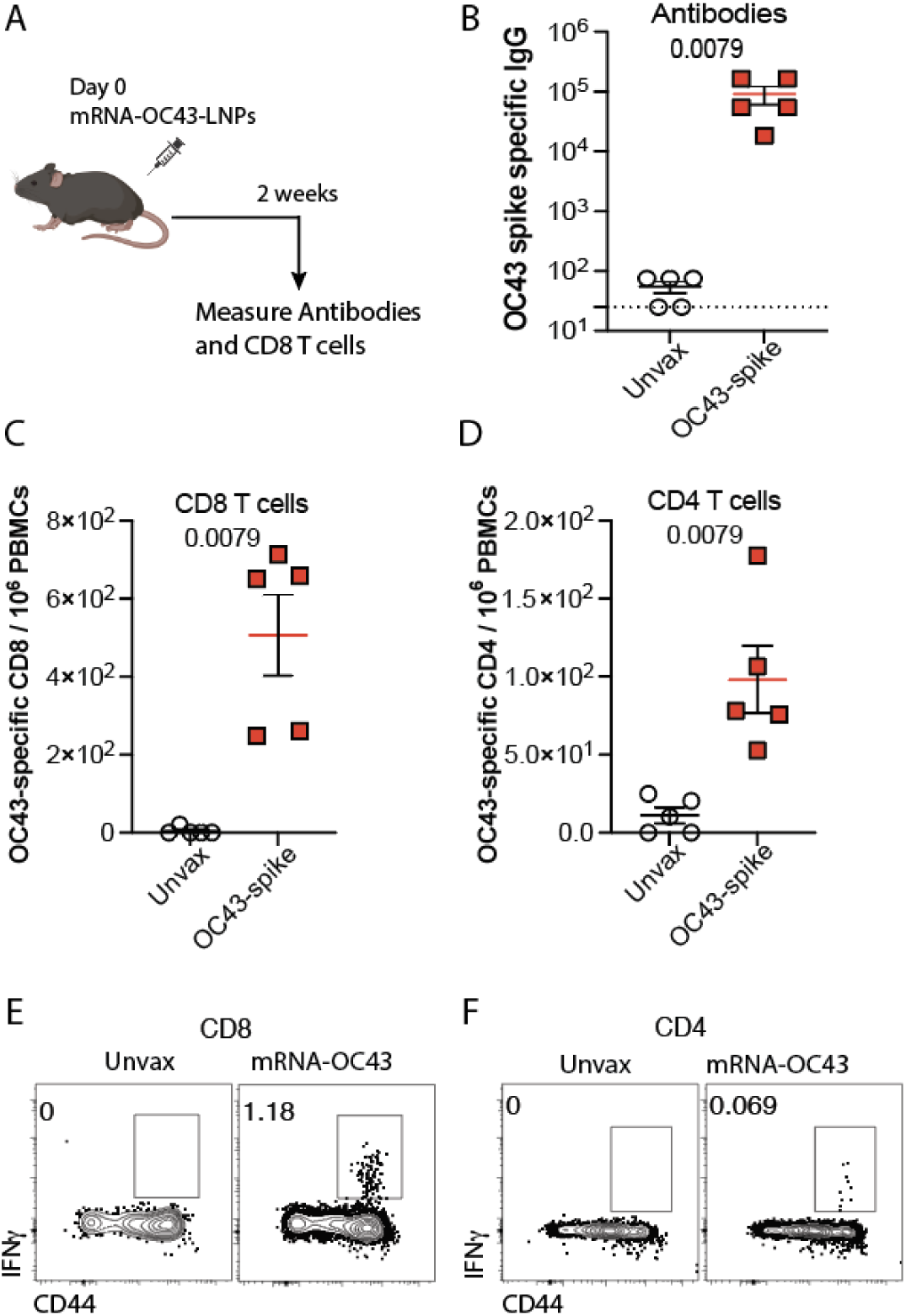
mRNA-OC43 elicits antibody and T cell responses. (A) Experimental outline representing the immunization scheme in C57BL/6 mice. Mice were immunized intramuscularly with 3 μg of OC43 spike mRNA vaccine and measured immune responses at day 15. (B) Summary of OC43-specific antibody responses in sera. (C to F) At day 15 after vaccination, PBMCs were incubated with overlapping OC43-spike peptide pools for 5 hours at 37^0^C in the presence of GolgiStop and GolgiPlug to detect OC43-specific CD8+ and CD4+ T cell responses. (C) Summary of OC43-specific CD8 T cells that express IFNγ in PBMCs. (D) Summary of OC43-specific CD4 T cells that express IFNγ in PBMCs. (E) Representative FACS plots showing the frequencies of OC43-specific CD8 T cells expressing IFNγ in PBMCs. (F) Representative FACS plots showing the frequencies of OC43-specific CD4 T cells expressing IFNγ in PBMCs. Data are from one experiment, n=5 mice per group. Indicated *P* values were determined by a nonparametric Mann-Whitney U test (unpaired t-test). Dashed lines indicate the limit of detection. Error bars represent SEM.

To examine vaccine efficacy, mice were challenged intranasally with 50 μL of OC43 neurovirulent (NV) strain, (1 × 10¹⁰ genome copies) at week 2 post-vaccination. Mice were monitored daily for any changes in clinical signs or symptoms, body mass, and mortality (Fig. 3A). Upon OC43 challenge, unvaccinated mice exhibited severe weight loss, severe clinical pathology and showed only 20% survival (Fig. 3B-3E). In contrast, mice that received the OC43 vaccine showed no weight loss and 100% survival, with no clinical signs of disease (Fig. 3F).

**Figure 3.**
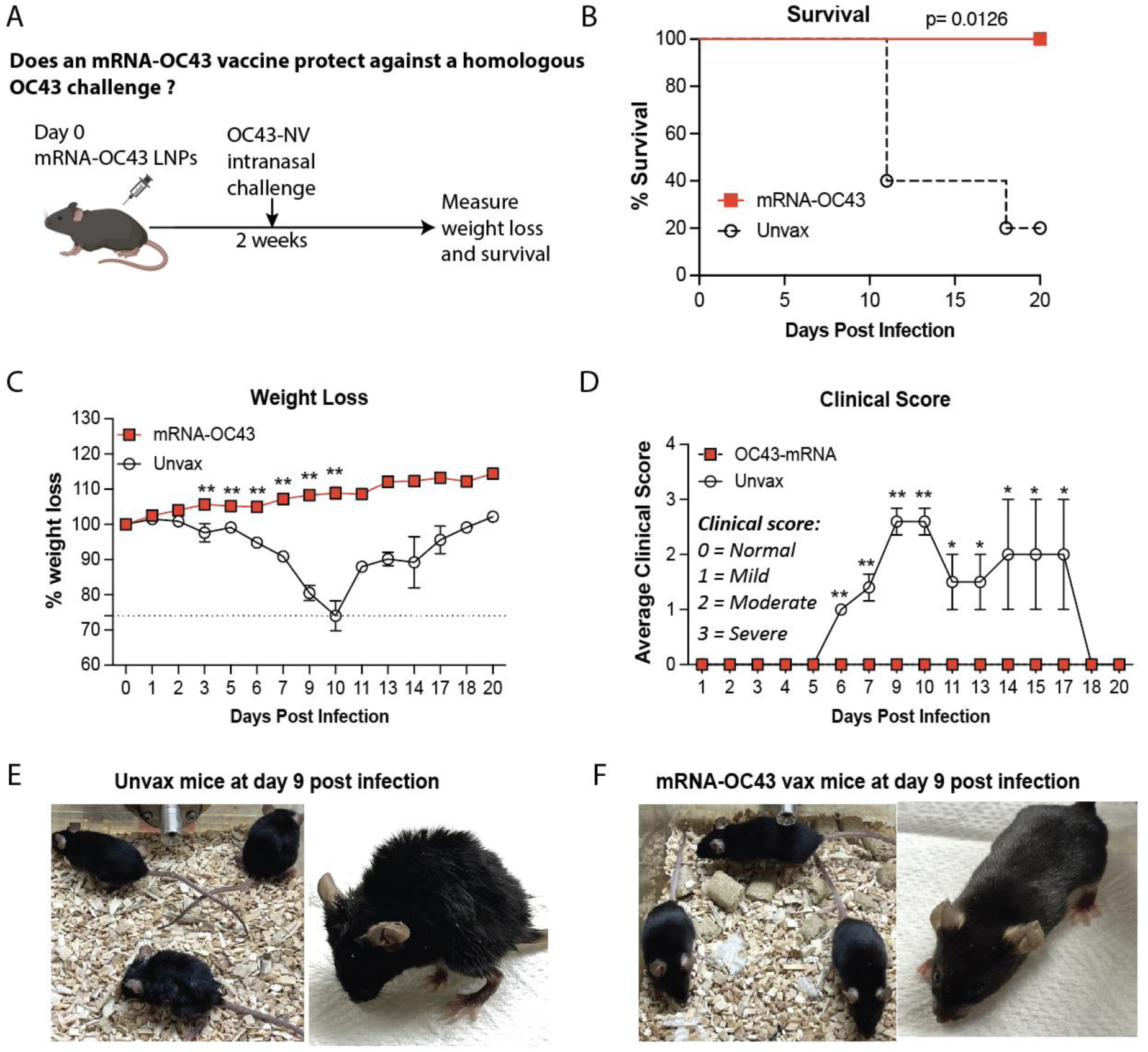
mRNA-OC43 vaccine protects mice from homologous OC43 infection. (A) experimental outline for evaluating whether OC43 mRNA vaccine protects mice against homologous OC43 infections. C57BL/6 mice were immunized with 3 μg of OC43-spike mRNA vaccine intramuscularly. Control mice received PBS. At 2 weeks following vaccination, 50 μL of OC43 neurovirulent strain (1 × 10¹⁰ genome copies) was intranasally inoculated into mice. Mice were monitored over 20 days for weight changes, mortality, and clinical signs of disease including hunched posture, ruffled fur, lethargy, and labored breathing. (B) Survival. (C) Weight loss. (D) Clinical score. Weight loss was calculated in terms of percent of original weight. Clinical scores were assigned on a scale of 0-3 in each category, where clinical scores 0, 1, 2, and 3 represent normal healthy, mild, moderate, and severe clinical signs/symptoms, respectively (see the details of the disease score in methods). Daily scores were averaged. (E) Representative images of unvaccinated mice at day 9 post OC43 infection. Mice showed severe clinical pathology including piloerection, puffy appearance, severely hunched posture, eyes completely closed, non-responsive, stopped eating and drinking, labored breathing with gasps. (F) Representative images of mRNA-OC43 immunized mice at day 9 post OC43 infection. Mice were active with smooth coat appearance. Clinical scores and weight were compared using multiple student’s t-tests with Holm-Sidak multiple comparison correction. Survival curves were compared using the log-rank Mantel-Cox comparison test. Significant differences compared to control are indicated \**p* < 0.05, \*\**p* < 0.01. Data are from one experiment, including 5 mice per group. Error bars represent SEM.

### Protective Efficacy of mRNA-OC43 following a heterologous coronavirus challenge

Further, we interrogated whether the mRNA-OC43 vaccine could cross-protect against another Embecovirus (MHV-A59). MHV-A59 is a well-studied mouse virus (11–13). OC43 and MHV-A59 share only ∼65% sequence identity in their spike proteins, rendering MHV-A59 a stringent challenge model to examine cross-protection by our mRNA-OC43 vaccine (Fig. 4). While MHV-A59 is not considered a significant threat to humans, it serves as a useful proof-of-principle model to evaluate the protective breadth of our OC43 vaccine.

**Figure 4.**
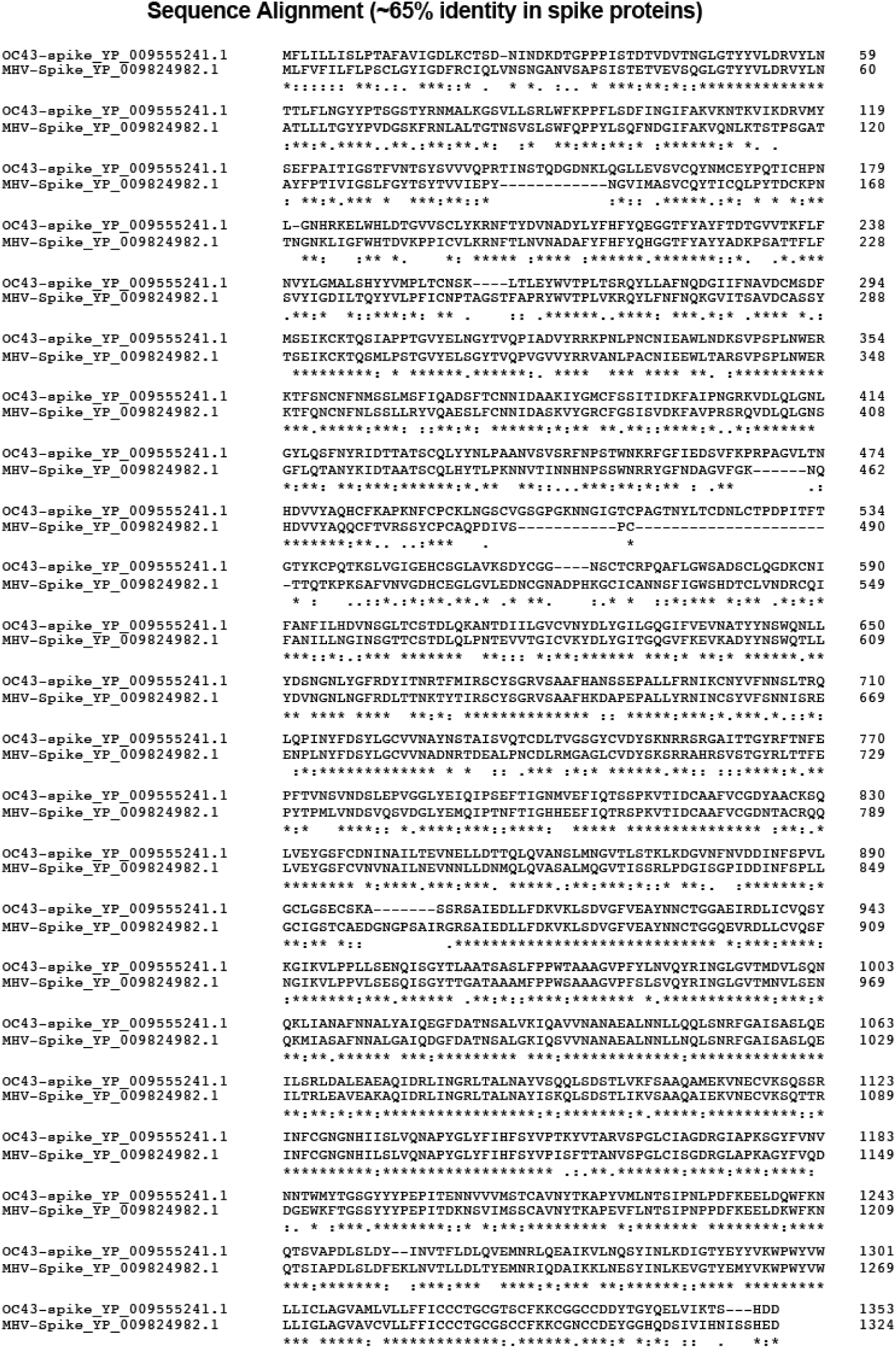
Sequence identity in spike glycoproteins of HCoV-OC43 and MHV-A59. Amino acid alignment of spike protein sequences of OC43 and MHV-A59 strains representing the 65 percent sequence identity in spike protein. Asterisks indicate identical residues; colons represent conserved changes, and blank spaces denote non-conserved substitutions.

To interrogate the cross-protective efficacy of OC43 vaccine, mice were immunized intramuscularly with 3 μg of the mRNA-OC43 vaccine. After 2 weeks post-vaccination, mice were challenged intraperitoneally with 2×10^6^ pfu of MHV-A59 (Fig. 5A). All mice experienced weight loss following infection, but the mice that were vaccinated with mRNA-OC43 vaccine exhibited significantly less weight loss compared to control (Fig. 5B). Further, the clinical signs were significantly milder in mRNA-OC43 vaccinated mice (Fig. 5C). All mice were sacrificed at day 3 post infection for assessing viral load in tissues. Importantly, the mRNA-OC43 vaccinated mice showed enhanced viral control in lung, brain, and liver (Fig. 5D-5F). These data suggest that the mRNA-OC43 vaccine provides cross-protection against a heterologous coronavirus infection with only a 65% antigen match.

**Figure 5.**
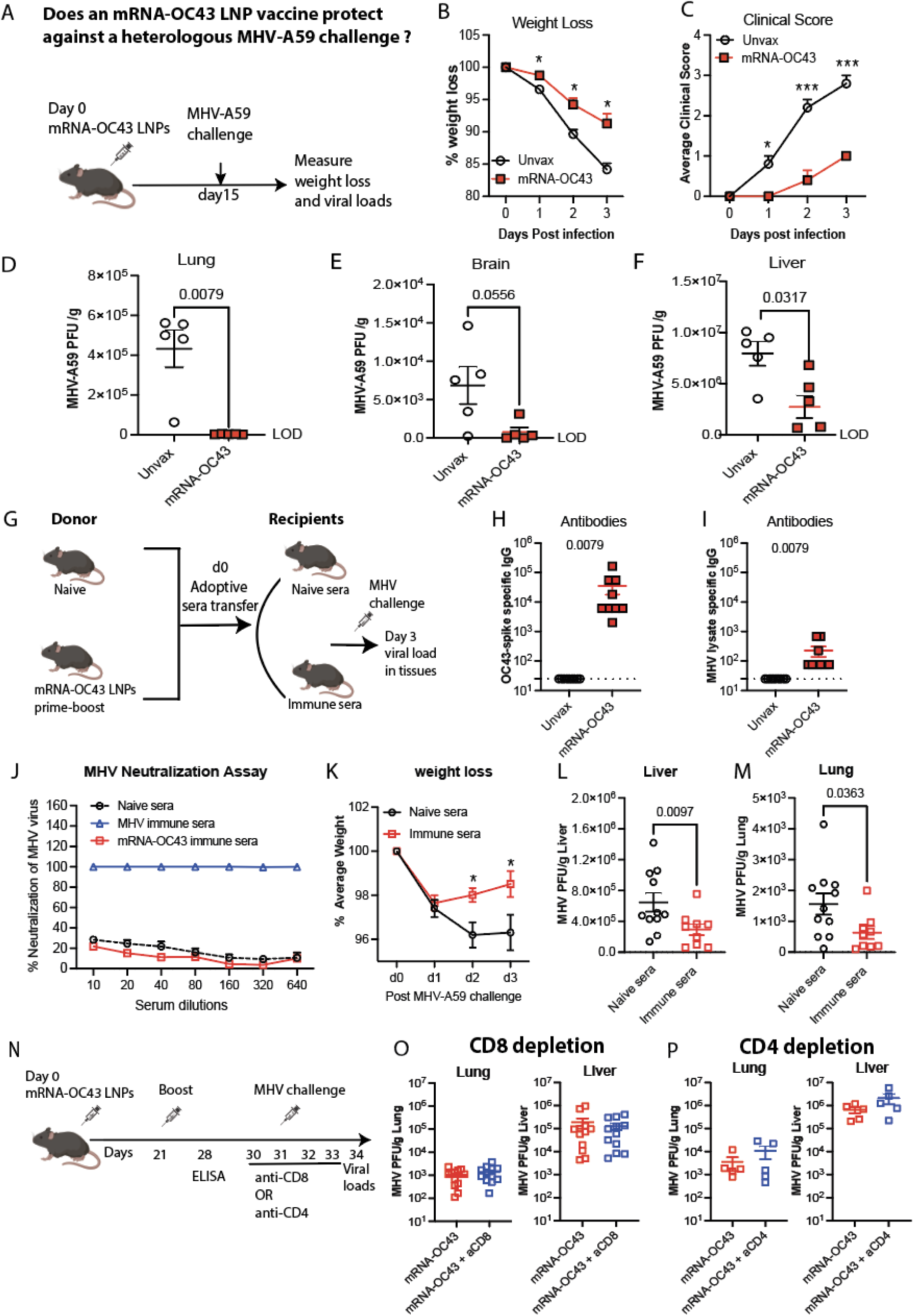
mRNA-OC43 vaccine protects mice against heterologous MHV-A59 infection. (A) Experimental outline for investigating the efficacy of OC43 mRNA vaccine in C57BL/6 mice upon MHV-A59 challenge. Mice were immunized with 3 μg of OC43-spike mRNA vaccine intramuscularly. Control mice received PBS. At 2 weeks following vaccination, all mice were inoculated with 500 μL of MHV-A59 (2×10⁶ PFU) via intraperitoneal route. Body weight and clinical signs of disease (hunched posture, ruffled fur, lethargy, and labored breathing) were measured daily over 3 days. Lung, brain, and liver were collected on day 3 post-infection to assess viral load by plaque assay. (B) Weight loss. Weight loss was calculated in terms of percent of original weight. (C) Clinical score. Clinical scores were assigned on a scale of 0-3 in each category, where clinical scores 0, 1, 2, and 3 represent normal healthy, mild, moderate, and severe clinical signs/symptoms, respectively (see methods for details). Daily scores were averaged. Clinical scores and weight were compared using multiple student’s t-tests with Holm-Sidak multiple comparison correction. (D) Viral load in the lung. LOD = 29 PFU/g. (E) Viral load in the brain. LOD = 11 PFU/g. (F) Viral load in the liver. LOD = 5 PFU/g. (G) Schematic layout of adoptive plasma transfer experiment: Donor mice were primed and boosted with 3 μg of mRNA-OC43 vaccine at 4 week-interval and harvested plasma on day 7 after boost. An 800 μL of pooled plasma per mouse was adoptively transferred into recipient mice via i.p. injection, followed by MHV-A59 challenge next day. Control mice received naïve plasma. Body weight and clinical signs were measured for three consecutive days. (H) Homologous OC43-specific antibody responses detected by ELISA in donor’s plasma at day 7 post-boost (or day 28 post-prime). (I) Heterologous cross-reactive antibody binding to MHV-A59 lysate in ELISA on day 7 post-boost plasma from donor mice. (J) MHV-A59 plaque reduction neutralization titer (PRNT) assay was performed using donor plasma collected at day 7 post-boost. Ten-fold serial dilutions of plasma from mRNA-OC43 immunized or naïve mice were incubated with MHV-GFP for 2 hours, then virus neutralized plasma mixture was added to L2 cells. The percentage of MHV neutralization was quantified by counting GFP-positive infected L2 cells using the Incucyte live-cell imaging system. Plasma from MHV-infected mice served as a positive control, achieving 100% neutralization. Naïve plasma served as a negative control. Viral load was assessed in tissues of recipient mice on day 3 post infection. (K) Weight loss. (L) Viral load in Liver. LOD = 5 PFU/g. (M) Viral load in Lung. LOD = 36 PFU/g. (N) Schematic of the T cell depletion experiments. Vaccinated and unvaccinated control mice were administered with CD8 or CD4 depleting antibodies for four consecutive days, starting one day before MHV-A59 challenge and continuing through day 3 post-infection. Viral loads in the lungs and liver were measured on day 3 post-infection. (O) Viral load in tissues of CD8-depleted mice. (P) Viral load in tissues of CD4-depleted mice. Indicated p values were determined by nonparametric Mann-Whitney U test (unpaired t-test). Significant differences compared to control are indicated \**p* < 0.05, \*\*\**p* < 0.001. Data are from one experiment (Panels A-F and P) or two experiments (Panels G-O), with n=5 mice per group. All data are shown. Dashed lines indicate the limit of detection. Error bars represent SEM.

### mRNA-OC43 elicited antibodies confer cross-protection against MHV-A59

Vaccine protection is typically mediated by humoral and cellular responses. To specifically assess the role of humoral protection, we performed a passive immunization study (Fig. 5G). First, we immunized C57BL/6 mice with the mRNA-OC43 vaccine on days 0 and 21 and then collected immune plasma on day 35. Prior to adoptive transfer, antibody titers specific to the OC43 spike antigen were confirmed in immune plasma using ELISA (Fig. 5H). These mRNA-OC43-immune plasma also exhibited cross-reactivity against a lysate of MHV-A59 infected 17CL-1 cells (Fig. 5I). However, these mRNA-OC43 immune plasma did not neutralize MHV-A59 in vitro, suggesting no or limited neutralization potential by plaque reduction neutralization titer assays (PRNT) (Fig. 5J). Each recipient mouse received 800 μL of pooled plasma (administered gradually over 96 h) via intraperitoneal injection. Control mice received plasma from naïve animals. On the following day, all recipient mice were challenged intraperitoneally with 2 × 10⁶ PFU of MHV-A59. Mice were monitored for three consecutive days for weight loss, and we determined viral load in tissues on day 3 after infection. Notably, mice that received immune plasma exhibited reduced weight loss and significantly lower viral loads in the lung (2.5-fold reduction) and liver (2.2-fold reduction) (Fig. 5K–5M), indicating that antibodies elicited by the mRNA-OC43 vaccine provide cross-protection against MHV-A59 infection, despite not exerting neutralizing function.

Next, we investigated whether T cells elicited by the mRNA-OC43 vaccine contribute to the control of MHV-A59 (Fig. 5N). To assess the role of CD8⁺ T cells, we depleted them at the time of infection in both immunized and control mice, using depleting antibodies. CD8⁺ T cell depletion had no observable effect on viral infection (Fig. 5O). Similarly, in separate experiments, we depleted CD4⁺ T cells to determine whether they impaired vaccine protection. CD4⁺ T cell depletion also did not impair vaccine protection (Fig. 5P). These findings suggest that T cells may not play a significant role in the cross-protection elicited by the mRNA-OC43 vaccine.

### mRNA-MHV vaccine confers heterologous protection against OC43

We have shown that an mRNA-OC43 vaccine confers heterologous protections against MHV. We also performed the “inverse” vaccination-challenge study. Mice were immunized intramuscularly with an mRNA-MHV vaccine followed by a lethal challenge with OC43 (Fig. 6A). On day 15 after vaccination, antibody and T cell responses were measured. As expected, the mRNA-MHV vaccine induced antibody responses against its matched antigen, MHV lysate (Fig. 6B). Interestingly, this mRNA-MHV vaccine also elicited cross-reactive antibody responses against other coronaviruses, including OC43, HKU1, and SARS-CoV-2 (Fig. 6C-6E). This vaccine not only elicited MHV-specific CD8⁺ T cell responses (K^b^S598) (Fig. 6F-6G), but also cross-reactive OC43-specific CD8^+^ T cell responses by ELISpot (Fig. 6H).

**Figure 6.**
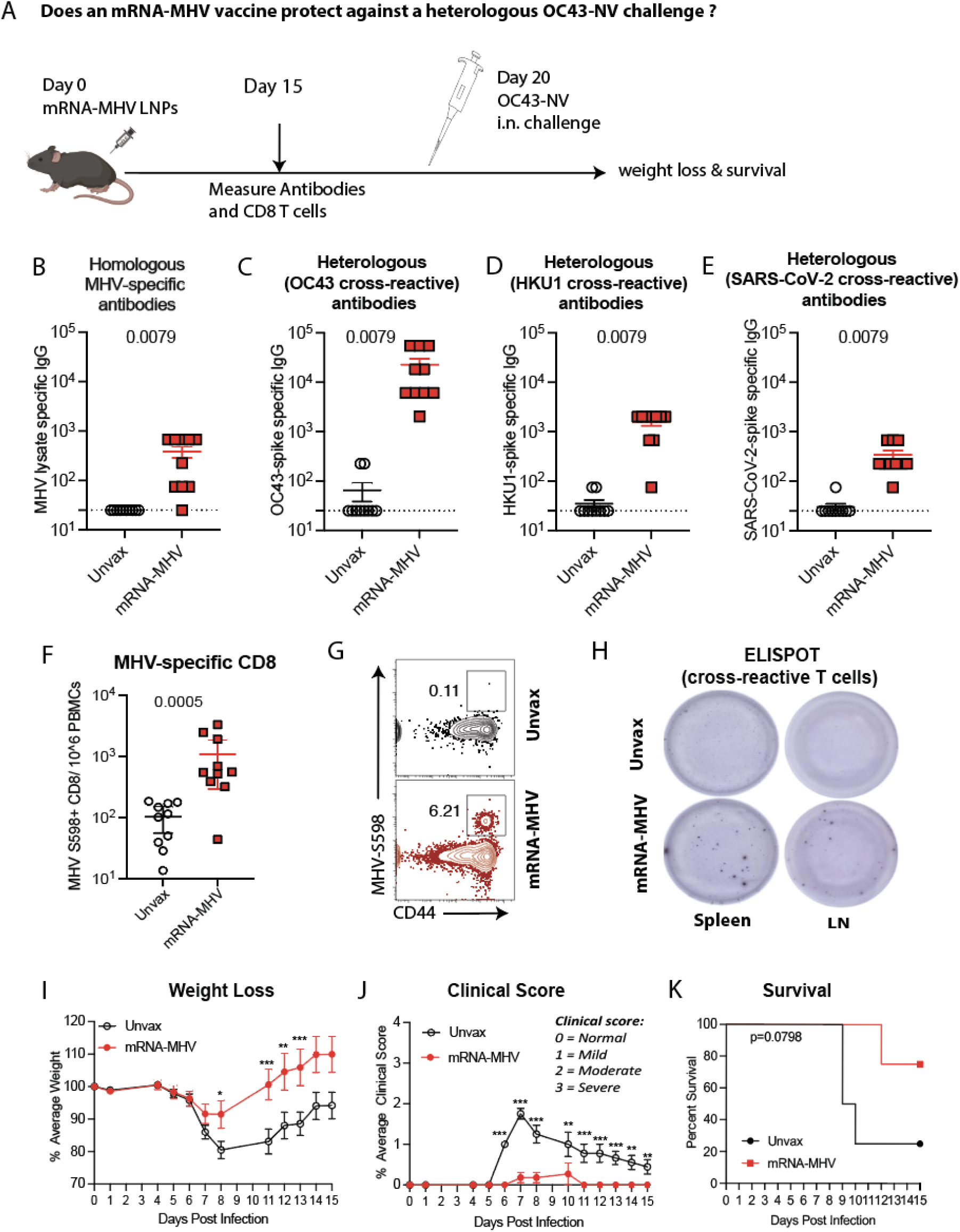
An mRNA-MHV-JHM provides cross- protection against heterologous OC43 infection. (A) Schematic representation of experiment investigating the cross-immune protection provided by mRNA-MHV vaccine against heterologous OC43 infection in C57BL/6 mice. Mice were primed with 5 μg of mRNA-MHV vaccine intramuscularly. Control mice received PBS. On day 20 after prime, mice were intranasally inoculated with 50 μL of OC43 (1 × 10¹⁰ genome copies). Body weight and clinical signs of disease (hunched posture, ruffled fur, lethargy, and labored breathing) were measured daily over two weeks. Antibody responses were tested at day 15 post vaccination by ELISA. (B) Summary of MHV-A59 lysate-specific antibody response. (C) Summary of OC43-spike-specific cross-reactive antibodies. (D) Summary of HKU1-spike-specific cross-reactive antibodies. (E) Summary of SARS-CoV-2-spike-specific cross-reactive antibodies. (F) Summary of MHV-spike (K^b^S598) specific CD8^+^ T cells in PBMCs at day 15 post vaccination. (G) Representative FACS plots of MHV-spike (K^b^S598) specific CD8^+^ T cells in PBMCs. (H) Representative wells of ELISpot plate showing the cross-reactive OC43-spike-specific T cells in spleen and draining lymph nodes. (I) Weight loss. Weight loss was calculated in terms of percent of original weight. (J) Clinical score. Clinical scores were assigned on a scale of 0-3 in each category, where clinical scores 0, 1, 2, and 3 represent normal healthy, mild, moderate, and severe clinical signs/symptoms, respectively (see methods for details). Daily scores were averaged. (K) Survival. Clinical scores and weight were compared using multiple student t-tests. Indicated p values were determined by nonparametric Mann-Whitney U test (unpaired t-test). Significant differences compared to control are indicated \**p* < 0.05, \*\**p* < 0.01, \*\*\**p* <0.001. Survival curves were compared using the log-rank Mantel-Cox comparison test. Data are from two experiments, with n=5 mice per group. All data are shown. Dashed lines indicate the limit of detection. Error bars represent SEM.

To evaluate the efficacy of the MHV vaccine, mice were intranasally challenged with OC43 neurovirulent strain three weeks after vaccination and then monitored for weight loss and clinical score. Following OC43 challenge, unvaccinated mice tended to show more significant weight loss and worse disease, compared to vaccinated mice (Fig. 6I–6J). There was also a pattern of improved survival with the mRNA-MHV vaccine relative to control, but the difference was not statistically significant (Fig. 6K). These results suggest that an mRNA-MHV vaccine confers partial protection against a distant OC43 challenge.

## DISCUSSION

There are four endemic human coronaviruses that typically cause mild respiratory infections. These include two Alphacoronaviruses (HCoV-229E and HCoV-NL63) and two Betacoronaviruses (HCoV-OC43 and HKU1), which are both part of the embecovirus sublineage. In this study, we focused on OC43, given the availability of a mouse model and the fact that it accounts for a great fraction of common cold coronavirus infections in humans (14, 15). OC43 belongs to the Betacoronavirus genus, alongside with SARS-CoV-2, SARS-CoV, and MERS-CoV, which were responsible for outbreaks in 2019, 2003, and 2012, respectively.

While previous studies have shown that coronavirus vaccines can generate cross-reactive antibodies against endemic coronaviruses, it remains unclear whether these antibodies are cross-protective in vivo (16–19). Building on this, we hypothesized that an mRNA vaccine targeting the human common cold coronavirus OC43, for which no effective vaccine currently exists, could also confer cross-protection against other coronaviruses. To test this hypothesis, we developed a novel mRNA vaccine encoding a stabilized OC43 spike protein and assessed its immunogenicity and protective efficacy in vivo. This mRNA-OC43 vaccine elicited adaptive immune responses to OC43 and conferred protection against OC43 infection, which was expected given that the vaccine antigen was matched to the challenge antigen. However, an interesting finding was that the mRNA-OC43 vaccine also provided cross-protection against MHV-A59, despite OC43 and MHV-A59 having only 65% identity in their spike proteins. Although the mRNA-OC43 vaccine induced antibodies that cross-reacted with MHV, these antibodies did not exhibit neutralizing activity. Adoptive plasma transfer experiments demonstrated that antibodies generated by the mRNA-OC43 vaccine protected mice against MHV-A59 infection. This result suggests that antibody-mediated protection occurred via effector functions and not via neutralization, supporting previous studies that demonstrate antiviral effects by non-neutralizing antibodies during viral infections (20–25). Interestingly, we did not observe a protective role for CD8⁺ or CD4⁺ T cells against MHV infection in our model, which contrasts with other reports demonstrating a critical role for virus-specific T cells in mediating cross-protection (26–29). One possible explanation is that the depleting antibodies used in our study may have effectively eliminated T cells in the blood but failed to do so efficiently in tissues like the lung or liver, potentially obscuring their contribution to cross-protection.

To further substantiate our findings, we immunized mice with an mRNA vaccine encoding the MHV spike protein and evaluated its protective efficacy against OC43 infection. This strategy also conferred cross-protection, as demonstrated by reduced weight loss, milder clinical symptoms, and increased survival rates in vaccinated animals. Notably, this mRNA-MHV vaccine elicited cross-binding antibody responses to multiple betacoronaviruses, including OC43, HKU1, and SARS-CoV-2, as well as cross-reactive T cell responses targeting the OC43 spike protein. These results suggest that vaccines targeting a single coronavirus strain can confer broad protection against other coronaviruses from the same subgenus. These data may be important for improving vaccine preparedness against circulating and emerging coronaviruses.

## ACKNOWLEDGEMENTS

This work was possible with grants from the National Institute on Drug Abuse (NIDA, DP2DA051912), Third Coast Centers for AIDS Research (CFAR), and the National Institute of Allergy and Infectious Diseases (NIAID, 1R56AI187084) to P.P.M. We thank Tom Gallagher (Loyola University); Susan R. Weiss (University of Pennsylvania) and Stanley Perlman / Noah Schuster (University of Iowa) for reagents and advice.

## AUTHOR CONTRIBUTIONS

P.P.M. and T.D. designed experiments. T.D. conducted prime-boost immunization and immunogenicity experiments in mice. S.L. helped in CD4 depletion experiment. T.D. made the mRNA vaccine. P.P.M., and T.D., wrote the paper, with feedback from all authors.

## MATERIALS AND METHODS

### Mice, and Immunizations

6-8-week-old C57BL/6 mice were used. Mice were purchased from Jackson laboratories (approximately half males and half females). Mice were immunized intramuscularly with mRNA-LNPs (made in-house) diluted in sterile PBS. Mice were housed at Northwestern University’s Center for Comparative Medicine (CCM). All mouse experiments were performed with the approval of the Northwestern University Institutional Animal Care and Use Committee (IACUC).

### Synthesis of modified mRNA

We synthesized mRNA vaccines encoding for the codon-optimized OC43 spike protein from HCoV-OC43 (Accession number AAA03055.1) and codon-optimized MHV-spike protein from MHV-JHM strain (Accession number YP_209233.1). For *in vitro* transcription of mRNA (IVT-mRNA), plasmid constructs were designed by incorporating codon-optimized immunogens (OC43-spike or MHV-spike), UTRs, and phase T7 RNA polymerase promoter and purchased from Genscript. The sequences of the 5′- and −3′-UTRs were identical to those used in a previous publication (25). Modified nucleotide pseudouridine-5’-triphosphate (ΨTP), along with canonical nucleotides ATP, CTP, and GTP (CellScript, Cat. No. ICTY110510), was used to synthesize nucleoside-modified IVT-mRNA from the plasmid construct. To enhance mRNA stability, an N7-methylguanosine cap (Cap 1, m⁷G) was added to the 5′ end, and a ∼150-nucleotide poly(A) tail was incorporated at the 3′ end using CellScript Capping and Tailing Kits (Cat. Nos. SCCS1710 and PAP5104H). The IVT-mRNA was purified via ammonium acetate precipitation and quantified using a NanoDrop ONE spectrophotometer (Thermo Scientific). To evaluate protein expression, purified mRNA was transfected into female human embryonic kidney (HEK) 293T cells using the TransIT-mRNA Transfection Kit (Mirus, Cat. No. MIR2250). Cell lysates from transfected HEK293T cells was analyzed by western blot to confirm spike protein expression. Following confirmation, the mRNA was encapsulated in lipid nanoparticles as described below.

### mRNA-LNP formulation

All purified mRNAs generated above were encapsulated into lipid nanoparticles using the NanoAssemblr Benchtop system (Precision NanoSystems) and confirmed to have similar encapsulation efficiency (∼95%). In brief, mRNA was diluted in 50 mM sodium acetate buffer, pH 5.0 to achieve a working concentration of 0.096 mg/mL (Cayman Chemical, Cat. No. 35425). An ethanolic lipid mixture was prepared using four lipids-SM-102, 1,2-distearoyl-sn-glycero-3-PC, Cholesterol, and DMG-PEG (Cayman, Cat. No. 35425) in a molar ratio of 50: 10: 38.5: 0.38. Subsequently, diluted mRNA in an aqueous phase and lipid mixture was run through a microfluidic laminar flow cartridge (NanoAssemblr Ignite NxGen, Cat. No. NIN0061). This was done by maintaining a nitrogen-to-phosphate (N/P) ratio of 4.0 (Lipid mix to mRNA ratio of 4), an RNA-to-lipid flow ratio of 3:1, and a total flow rate of 12 mL/min to generate mRNA–lipid nanoparticles (mRNA-LNPs). The resulting mRNA-LNPs were concentrated and purified using an Amicon Ultra-15 filtration unit and a 0.2 µm Acrodisc filter. Encapsulation efficiency and the concentration of encapsulated mRNA were determined using the Quant-iT RiboGreen RNA Assay Kit (Invitrogen, Cat. No. R11490).

### Reagents, Flow cytometry, and Equipment

To determine the T cell responses in the blood and spleen, single-cell suspensions of PBMCs and spleen were prepared as described previously (30). Dead cells were gated out using Live/Dead fixable dead cell stain (Invitrogen). The CD8 and CD4 responses specific to the OC43 spike were measured by stimulating splenocytes with the OC43 spike peptide pools (NR-53728, BEI) in intracellular cytokine staining (ICS). Biotinylated MHC class I monomer (K^b^S598, sequence RCQIFANI) was used for detecting MHV spike-specific CD8 T cells and was obtained from the NIH tetramer facility at Emory University. Cells were stained with fluorescently labeled antibodies against anti-mouse CD8α (53-6.7 on PerCP-Cy5.5), anti-mouse CD4 (RM4-5 FITC), anti-mouse CD44 (IM7 on Pacific Blue), anti-mouse IFNγ (XMG1.2 on APC). Fluorescently labeled antibodies were purchased from BD Pharmingen, except for anti-CD44 (which was from Biolegend). Dead cells were gated out using LIVE/ DEAD fixable dead cell stain (Invitrogen). Flow cytometry samples were acquired with a Becton Dickinson Canto II or an LSRII and analyzed using FlowJo v10 (Treestar). The following reagent was obtained through BEI Resources, NIAID, NIH: Peptide Array, Human Coronavirus OC43 Spike (S) Glycoprotein, NR-53728.

### OC43 spike, HKU1 spike, SARS-CoV-2 spike and MHV lysate-specific ELISA

Binding antibody titers were measured using ELISA as described previously (31, 32). In brief, 96-well flat bottom plates MaxiSorp (Thermo Scientific) were coated with 0.1 μg/well of the respective spike protein for 24 hr at 4^0^C. For detection of MHV-specific antibody responses, MHV-A59 lysates were used as coating antigens (incubated for 48 hr at room temperature). Plates were washed with PBS + 0.05% Tween-20. Blocking was performed for 4 hr at room temperature with 200 μL of PBS + 0.05% Tween-20 + bovine serum albumin. 6μL of sera were added to 144 μL of blocking solution in the first column of the plate, 1:3 serial dilutions were performed until row 12 for each sample, and plates were incubated for 60 minutes at room temperature. Plates were washed three times followed by the addition of goat anti-mouse IgG horseradish peroxidase-conjugated (Southern Biotech) diluted in blocking solution (1:5000), at 100 μL/well and incubated for 60 minutes at room temperature. Plates were washed three times and 100 μL /well of Sure Blue substrate (Sera Care) was added for approximately 8 minutes. The reaction was stopped using 100 μL/well of KPL TMB stop solution (Sera Care). Absorbance was measured at 450 nm using Spectramax Plus 384 (Molecular Devices). OC43 spike protein was produced in-house using a mammalian expression vector obtained from Addgene (Cat. No. 166015).

### Propagation and determination of OC43-NV titers

OC43-NV stocks were propagated in 1-2 days old suckling neonates from C57BL/6 mice using a protocol from a prior paper (33). In brief, ten 1–2-day-old mice were inoculated intracerebrally with 10 μL of brain homogenates infected with OC43-NV (kind gift from Dr. Stanley Perlman’s laboratory). After two days, whole brains were collected from the neonates and homogenized in 2 mL of sterile PBS. The lysates were clarified by centrifugation at 2000 rpm for 10 minutes, as previously described (34) and small aliquots of the supernatant were stored in a −80°C. For adult mouse challenges, the above viral stock was diluted 10-fold and 50 μL was administered intranasally. Viral load in brain lysates was quantified by quantitative real-time RT-PCR targeting the OC43 nucleocapsid gene, using TaqMan chemistry as previously described (19).

### OC43-NV challenge and disease severity score

On day 15 post-vaccination, mice were challenged intranasally with 50 μL of OC43-NV stock (1 × 10¹⁰ genome copies), administered as 25 μL per nostril. All mice were monitored for weight loss and clinical severity over the course of three weeks. Disease severity was measured in terms of clinical scores ranging from 0-3, defining the body posture, appearance of fur, eye secretions or closure, animal activity, lethargy, body temperature, and neurological symptoms (35, 36). The highest score was represented by severe disease status counting piloerection, puffy appearance, non-responsive and stationary even when provoked, severely hunched posture, completely closed eyes, stopped eating/drinking, rapid or labored breathing with gasps, cold body temperature, shivering, showing no response upon stimuli, and neurological symptoms. Score 2 was defined as moderate disease including moderately hunched posture, majority of fur on back is ruffled, active only when provoked, stationary, no response to auditory or slowed response to touch, eyes half-closed, potential eye secretions, lethargy, less active, and consistently labored breathing. The mild disease was scored as one showing mildly hunched, slightly ruffled fur, active, avoids standing upright, slowed response to auditory/touch stimuli. The normal active animal with smooth coat was scored as zero.

### MHV propagation and quantification

Seed stock of MHV-A59 was obtained from ATCC. The virus was propagated in 17CL-1 cells and tittered on L2 cells (kind gift from Dr. Susan R. Weiss). 17CL-1 and L2 cells were passaged in DMEM supplemented with 10% fetal bovine serum (FBS), 1% L-Glutamine and 1% Penicillin/Streptomycin. For virus propagation, 17CL-1 cells were inoculated at a low multiplicity of infection (MOI) of 0.1 in 1% DMEM. After 72 hours of incubation, the supernatant was collected and clarified by centrifugation at 2,000 rpm for 10 minutes. The titer of the viral stock was determined by plaque assay using L2 cell monolayers. To determine the viral titer in infected tissues, the lung, liver and brain were collected on day 3 after the challenge and stored immediately in a −80^0^C until processing. For plaque assay, 1 × 10^6^ L2 cells per well were seeded in 6-well plates in 10% DMEM medium. After 24 hours, when the monolayer was 90-100% confluent, tissue samples were thawed in water bath at 37^0^C and processed to assess viral titer. The tissues were homogenized using a standard TissueRuptur homogenizer, and 10-fold serial dilutions of the homogenized samples were prepared in 1% DMEM and applied dropwise onto the cell monolayer. The 6-well plates were placed in a 37°C, 5% CO_2_ incubator for 1 hour and manually rocked every 10 minutes. After 1 hour of incubation, a 1:1 agarose and 2×199 overlay was added to the monolayer, and plates were incubated at 37°C, 5% CO_2_ for 48 hours. After 48 hours, the overlay was removed, the cells were stained with 0.1% crystal violet, and plaques were counted.

### Adoptive plasma transfers

C57BL/6 donor mice were immunized with two doses of the mRNA-OC43 vaccine at a 3-week interval. Seven days after the final dose, OC43 spike-specific antibody responses were confirmed by ELISA, and plasma from the immunized mice was pooled. A total of 800 μL of this pooled plasma was adoptively transferred into C57BL/6 recipient mice via the intraperitoneal route. Control mice received plasma from naïve donors. The following day, all mice were infected intraperitoneally with 2 × 10⁶ PFU of MHV-A59. On day 3 post-infection, tissues were harvested and homogenized using a TissueRuptor Homogenizer (QIAGEN). Viral loads were quantified by plaque assay, as described above.

### Antibody treatments for CD4 and CD8 depletion

All antibodies used for in vivo treatments were purchased from BioXCell or Leinco, diluted in sterile PBS, and administered via intraperitoneal (i.p.) injection. CD4⁺ T cell-depleting antibody (GK1.5) and CD8⁺ T cell-depleting antibody (2.43) were given at a dose of 200 µg daily, starting one day before infection and continuing through day three post-infection. IgG isotype controls were included in all experiments as controls.

### Detection of IFN-γ-Producing T Cell Responses via ELISpot

To detect IFN-γ-producing antigen-specific T cells, 96-well ELISpot plates (Millipore, Burlington, MA) were coated with anti-mouse IFN-γ monoclonal antibody (clone AN-18, BioLegend 517902) at 5 μg/mL and incubated overnight at 4 °C. The following day, plates were washed twice with 200 μL/well of sterile 1× PBS and blocked with 200 μL/well of RPMI medium supplemented with 10% FBS, 1% L-glutamine, and 1% Pen/Strep for 2 hours at 37 °C in a CO₂ incubator. Single-cell suspensions from spleen or lymph nodes were prepared at 2.5 × 10⁶ cells/mL in supplemented RPMI. After blocking, media was discarded, and plates were seeded with 2.5 × 10⁵ cells/well and stimulated using 4 μg/mL of an OC43-spike peptide pools. Cells were incubated for 18–20 hours at 37 °C in a CO₂ incubator. Cells from naïve mice served as negative controls. For positive controls, cells were treated with either 10 ng/mL phorbol 12-myristate 13-acetate (PMA, Sigma) plus 500 ng/mL ionomycin (Sigma), or with anti-mouse CD3 and CD28 antibodies (1 μg/well). Unstimulated controls included cells incubated with media alone. After 18-20 hours of incubation, cells were discarded, and plates were washed five times with wash buffer (1× PBS + 0.05% Tween-20). Plates were then incubated for 90 minutes with biotinylated anti-IFN-γ antibody (clone R4-6A2, BioLegend 505704) at 0.5 μg/mL diluted in PBS with 10% FBS. After washing, streptavidin–alkaline phosphatase (Bio-Rad 170-6432), diluted 1:1000 in 10% PBS, was added and incubated for 45 minutes. Plates were washed with wash buffer and developed using substrate (Bio-Rad 170-6432) for 8 minutes. The reaction was stopped by rinsing plates with running water. Spot-forming cells (SFCs) were analyzed using an ImmunoSpot Image Analyzer (Cleveland, USA).

### Statistical analysis

Statistical tests used are indicated on each figure legend. Dashed lines in data figures represent the limit of detection. Statistical significance was established at a *p* value of 0.05 or less and was generally assessed by two tailed unpaired student’s t tests (Mann-Whitney *U* test), unless indicated otherwise in figure legends. Data were analyzed using Prism version 10 (GraphPad Software).

## REFERENCES

1. El-Sahly HM, Atmar RL, Glezen WP, Greenberg SB. 2000. Spectrum of clinical illness in hospitalized patients with “common cold” virus infections. Clin Infect Dis 31:96–100.

2. Vabret A, Mourez T, Gouarin S, Petitjean J, Freymuth F. 2003. An outbreak of coronavirus OC43 respiratory infection in Normandy, France. Clin Infect Dis 36:985–9.

3. Falsey AR, Walsh EE, Hayden FG. 2002. Rhinovirus and coronavirus infection-associated hospitalizations among older adults. J Infect Dis 185:1338–41.

4. Pene F, Merlat A, Vabret A, Rozenberg F, Buzyn A, Dreyfus F, Cariou A, Freymuth F, Lebon P. 2003. Coronavirus 229E-related pneumonia in immunocompromised patients. Clin Infect Dis 37:929–32.

5. van der Hoek L. 2007. Human coronaviruses: what do they cause? Antivir Ther 12:651–8.

6. McIntosh K, Chao RK, Krause HE, Wasil R, Mocega HE, Mufson MA. 1974. Coronavirus infection in acute lower respiratory tract disease of infants. J Infect Dis 130:502–7.

7. Walsh EE, Shin JH, Falsey AR. 2013. Clinical impact of human coronaviruses 229E and OC43 infection in diverse adult populations. J Infect Dis 208:1634–42.

8. Gaunt ER, Hardie A, Claas EC, Simmonds P, Templeton KE. 2010. Epidemiology and clinical presentations of the four human coronaviruses 229E, HKU1, NL63, and OC43 detected over 3 years using a novel multiplex real-time PCR method. J Clin Microbiol 48:2940–7.

9. Paules CI, Marston HD, Fauci AS. 2020. Coronavirus Infections—More Than Just the Common Cold. JAMA 323:707–708.

10. Choi WI, Kim IB, Park SJ, Ha EH, Lee CW. 2021. Comparison of the clinical characteristics and mortality of adults infected with human coronaviruses 229E and OC43. Sci Rep 11:4499.

11. Lavi E, Gilden DH, Wroblewska Z, Rorke LB, Weiss SR. 1984. Experimental demyelination produced by the A59 strain of mouse hepatitis virus. Neurology 34:597–603.

12. Yang Z, Du J, Chen G, Zhao J, Yang X, Su L, Cheng G, Tang H. 2014. Coronavirus MHV-A59 infects the lung and causes severe pneumonia in C57BL/6 mice. Virol Sin 29:393–402.

13. Boscarino JA, Logan HL, Lacny JJ, Gallagher TM. 2008. Envelope protein palmitoylations are crucial for murine coronavirus assembly. J Virol 82:2989–99.

14. Zeng ZQ, Chen DH, Tan WP, Qiu SY, Xu D, Liang HX, Chen MX, Li X, Lin ZS, Liu WK, Zhou R. 2018. Epidemiology and clinical characteristics of human coronaviruses OC43, 229E, NL63, and HKU1: a study of hospitalized children with acute respiratory tract infection in Guangzhou, China. Eur J Clin Microbiol Infect Dis 37:363–369.

15. Zhou Z, Qiu Y, Ge X. 2021. The taxonomy, host range and pathogenicity of coronaviruses and other viruses in the Nidovirales order. Anim Dis 1:5.

16. Anderson EM, Goodwin EC, Verma A, Arevalo CP, Bolton MJ, Weirick ME, Gouma S, McAllister CM, Christensen SR, Weaver J, Hicks P, Manzoni TB, Oniyide O, Ramage H, Mathew D, Baxter AE, Oldridge DA, Greenplate AR, Wu JE, Alanio C, D’Andrea K, Kuthuru O, Dougherty J, Pattekar A, Kim J, Han N, Apostolidis SA, Huang AC, Vella LA, Kuri-Cervantes L, Pampena MB, Betts MR, Wherry EJ, Meyer NJ, Cherry S, Bates P, Rader DJ, Hensley SE. 2021. Seasonal human coronavirus antibodies are boosted upon SARS-CoV-2 infection but not associated with protection. Cell 184:1858–1864.e10.

17. Saunders KO, Lee E, Parks R, Martinez DR, Li D, Chen H, Edwards RJ, Gobeil S, Barr M, Mansouri K, Alam SM, Sutherland LL, Cai F, Sanzone AM, Berry M, Manne K, Bock KW, Minai M, Nagata BM, Kapingidza AB, Azoitei M, Tse LV, Scobey TD, Spreng RL, Rountree RW, DeMarco CT, Denny TN, Woods CW, Petzold EW, Tang J, Oguin TH, 3rd, Sempowski GD, Gagne M, Douek DC, Tomai MA, Fox CB, Seder R, Wiehe K, Weissman D, Pardi N, Golding H, Khurana S, Acharya P, Andersen H, Lewis MG, Moore IN, Montefiori DC, Baric RS, Haynes BF. 2021. Neutralizing antibody vaccine for pandemic and pre-emergent coronaviruses. Nature 594:553–559.

18. Jacob-Dolan C, Feldman J, McMahan K, Yu J, Zahn R, Wegmann F, Schuitemaker H, Schmidt AG, Barouch DH. 2021. Coronavirus-Specific Antibody Cross Reactivity in Rhesus Macaques Following SARS-CoV-2 Vaccination and Infection. J Virol 95.

19. Dangi T, Palacio N, Sanchez S, Park M, Class J, Visvabharathy L, Ciucci T, Koralnik IJ, Richner JM, Penaloza-MacMaster P. 2021. Cross-protective immunity following coronavirus vaccination and coronavirus infection. J Clin Invest 131.

20. Clark JJ, Hoxie I, Adelsberg DC, Sapse IA, Andreata-Santos R, Yong JS, Amanat F, Tcheou J, Raskin A, Singh G, González-Domínguez I, Edgar JE, Bournazos S, Sun W, Carreño JM, Simon V, Ellebedy AH, Bajic G, Krammer F. 2024. Protective effect and molecular mechanisms of human non-neutralizing cross-reactive spike antibodies elicited by SARS-CoV-2 mRNA vaccination. Cell Rep 43:114922.

21. Izadi A, Nordenfelt P. 2024. Protective non-neutralizing SARS-CoV-2 monoclonal antibodies. Trends Immunol 45:609–624.

22. Asthagiri Arunkumar G, Ioannou A, Wohlbold TJ, Meade P, Aslam S, Amanat F, Ayllon J, García-Sastre A, Krammer F. 2019. Broadly Cross-Reactive, Nonneutralizing Antibodies against Influenza B Virus Hemagglutinin Demonstrate Effector Function-Dependent Protection against Lethal Viral Challenge in Mice. J Virol 93.

23. Tauzin A, Nayrac M, Benlarbi M, Gong SY, Gasser R, Beaudoin-Bussières G, Brassard N, Laumaea A, Vézina D, Prévost J, Anand SP, Bourassa C, Gendron-Lepage G, Medjahed H, Goyette G, Niessl J, Tastet O, Gokool L, Morrisseau C, Arlotto P, Stamatatos L, McGuire AT, Larochelle C, Uchil P, Lu M, Mothes W, De Serres G, Moreira S, Roger M, Richard J, Martel-Laferrière V, Duerr R, Tremblay C, Kaufmann DE, Finzi A. 2021. A single dose of the SARS-CoV-2 vaccine BNT162b2 elicits Fc-mediated antibody effector functions and T cell responses. Cell Host Microbe 29:1137–1150.e6.

24. Vogt MR, Dowd KA, Engle M, Tesh RB, Johnson S, Pierson TC, Diamond MS. 2011. Poorly neutralizing cross-reactive antibodies against the fusion loop of West Nile virus envelope protein protect in vivo via Fcgamma receptor and complement-dependent effector mechanisms. J Virol 85:11567–80.

25. Dangi T, Sanchez S, Class J, Richner M, Visvabharathy L, Chung YR, Bentley K, Stanton RJ, Koralnik IJ, Richner JM, Penaloza-MacMaster P. 2022. Improved control of SARS-CoV-2 by treatment with a nucleocapsid-specific monoclonal antibody. J Clin Invest 132.

26. Dos Santos Alves RP, Timis J, Miller R, Valentine K, Pinto PBA, Gonzalez A, Regla-Nava JA, Maule E, Nguyen MN, Shafee N, Landeras-Bueno S, Olmedillas E, Laffey B, Dobaczewska K, Mikulski Z, McArdle S, Leist SR, Kim K, Baric RS, Ollmann Saphire E, Elong Ngono A, Shresta S. 2024. Human coronavirus OC43-elicited CD4(+) T cells protect against SARS-CoV-2 in HLA transgenic mice. Nat Commun 15:787.

27. Humbert M, Olofsson A, Wullimann D, Niessl J, Hodcroft EB, Cai C, Gao Y, Sohlberg E, Dyrdak R, Mikaeloff F, Neogi U, Albert J, Malmberg KJ, Lund-Johansen F, Aleman S, Björkhem-Bergman L, Jenmalm MC, Ljunggren HG, Buggert M, Karlsson AC. 2023. Functional SARS-CoV-2 cross-reactive CD4(+) T cells established in early childhood decline with age. Proc Natl Acad Sci U S A 120:e2220320120.

28. Mallajosyula V, Ganjavi C, Chakraborty S, McSween AM, Pavlovitch-Bedzyk AJ, Wilhelmy J, Nau A, Manohar M, Nadeau KC, Davis MM. 2021. CD8(+) T cells specific for conserved coronavirus epitopes correlate with milder disease in COVID-19 patients. Sci Immunol 6.

29. Sagar M, Reifler K, Rossi M, Miller NS, Sinha P, White LF, Mizgerd JP. 2021. Recent endemic coronavirus infection is associated with less-severe COVID-19. J Clin Invest 131.

30. Sanchez S, Palacio N, Dangi T, Ciucci T, Penaloza-MacMaster P. 2021. Fractionating a COVID-19 Ad5-vectored vaccine improves virus-specific immunity. Sci Immunol 6:eabi8635.

31. Dangi T, Sanchez S, Lew MH, Visvabharathy L, Richner J, Koralnik IJ, Penaloza-MacMaster P. 2022. Pre-existing immunity modulates responses to mRNA boosters. bioRxiv doi:10.1101/2022.06.27.497248.

32. Sanchez S, Dangi T, Awakoaiye B, Lew MH, Irani N, Fourati S, Penaloza-MacMaster P. 2024. Delayed reinforcement of costimulation improves the efficacy of mRNA vaccines in mice. J Clin Invest 134.

33. Butler N, Pewe L, Trandem K, Perlman S. 2006. Murine encephalitis caused by HCoV-OC43, a human coronavirus with broad species specificity, is partly immune-mediated. Virology 347:410–21.

34. Xie P, Fang Y, Baloch Z, Yu H, Zhao Z, Li R, Zhang T, Li R, Zhao J, Yang Z, Dong S, Xia X. 2022. A Mouse-Adapted Model of HCoV-OC43 and Its Usage to the Evaluation of Antiviral Drugs. Front Microbiol 13:845269.

35. Gonzalez AJ, Ijezie EC, Balemba OB, Miura TA. 2018. Attenuation of Influenza A Virus Disease Severity by Viral Coinfection in a Mouse Model. J Virol 92.

36. Couto J, Gonçalves R, Lamas S, Saraiva M. 2023. Protocol for infecting and monitoring susceptible k18-hACE2 mice with SARS-CoV-2. STAR Protoc 4:102303.

